# Low-Cost, Modular Modification to a Desktop 3D Printer for General Purpose Gel/Paste Extrusion & Direct Ink Writing

**DOI:** 10.1101/2021.03.10.434735

**Authors:** D. J. Leech, S. Lightfoot, D. Huson, A. Stratakos

## Abstract

We propose a design for a simple paste extruder modification that can be used for the selective deposition and patterning of gels and pastes, using a desktop 3D printer as the primary platform. This technology has found use with a variety of materials in seemingly disparate fields, including the printing of ceramics, food and biological materials, each with a variety of material-specific solutions to enhance printability. However, we focus on a syringe-pump driven system that is simple, low-cost, modular, easily assembled and highly modifiable with a low barrier of entry in order to maximise the generalisability and range of printable materials.

## 1 Introduction

The precision positioning and selective deposition possible with extrusion based 3D printing platforms has relied heavily on the use of known and very specific filament formulations, such as PLA [1] and ABS [2], that allow for robust and versatile designs. However, many recent subsets of the field of additive manufacture have focused on the design and deposition principles of the process, whilst varying the printing materials to include biomaterials [3, 4, 5], glass [6, 7], a variety of other polymers [8, 9], metals [10, 11], concrete [12, 13], food [14, 15], and ceramics [16, 17].

The latter two examples are particularly interesting in their overlap with fused deposition modelling (FDM) printing techniques, due to their long history with other extrusion-based techniques, particularly within an industrial setting. However, while these examples historically work with relatively large amounts of material, bioprinting for instance tends toward small feature sizes and low material content, focusing on the building of structure from traditionally soft materials with high precision and maintaining any cell content within the ink [3]. As such, components of extrusion and extrusion-centred design and manufacture arise in a broad range of fields and approaches, in which materials in a reservoir are shaped, formed and built whilst under pressure caused by a die or nozzle. The name of this process can vary depending on the field, however we use material extrusion [18] or direct ink writing [19] as a ‘catch-all’ term to describe both the extrusion technique and the selective deposition enabled by modern 3D printing platforms. In terms of printable materials, we focus on a range of pastes or gels -generic terms to describe any viscous or relatively slow-flowing material.

Similar material generic extruders, utilise the selective XYZ positioning of a 3D printing platform, guiding a nozzle that is fed by a reservoir. The primary differences between these systems being:

- The method of feeding the material from the reservoir to the nozzle.
- Whether the reservoir system is mounted directly to the nozzle or fed via a connecting tube.
- The dimensions used (reservoir size, nozzle diameter, connector dimensions etc.)
- Any material specific optional upgrades (heating, cooling, enclosures etc.)

As mentioned, these systems allow for a large range of materials to be printed, however they are often subject to the same issues, including slow printing speeds, small reservoir sizes, requirements provided by the use of specific printer platforms, poor stability of printed artefacts, extensive stringing, single material extrusion and uneven extrusion [23]. Multiple designs have been proposed in recent years to tackle specific issues within the paste extruder system and low-cost bioprinters with varying degrees of complexity and printer specific additions, including Ref. [20], [21], [22], [23], [25] [26], [27], [28], [29], [30] and [31].

We propose a low-cost, generic and modular addition to a standard desktop 3D printer that allows for the deposition of a variety of gels and pastes -build files, scripts and G-Code can be found on the Github Paste Extruder project page. This design focuses on the need for a low-cost and low barrier of entry paste extruder add-on that is compatible with a variety of 3D printer platforms, achieved by containing the majority of the device separate from the printer, and placing very few restrictions on the nozzle, reservoir and connector dimensions. This generic approach to the dimensions allows for the possibility of scaling and means that both larger scale extrusion (up to cm width) and microextrusion should both be possible on this platform. Additionally, the simplicity of the design allows this modification to be made in the home, as well as in the lab, requiring no specialist assembly equipment beyond an FDM 3D printer.

We additionally explore the material properties of a variety of inks and design a handful of test procedures that allow, without extensive outside equipment, for judgements of how to tune the printability of a particular ink recipe. Every material has a unique response and therefore may require differing formulations, temperature, pressure and tweaking of other parameters in order to make reproducible and controlled prints to the desired standard. We instead intend to produce a general guide for a generic paste extruder system, highlighting possible and common routes of exploration that may be used to enhance preliminary results for a variety of ink formulations.

## 2 Paste Extruder Design

The paste extruder setup is intended to be modular and malleable, allowing for each component to be traded for other similar designs, depending on the needs of the user. This also means that the system can be quickly converted back into a traditional 3D printer platform with minimal effort. We use a simple setup comprising of a desktop 3D printer, syringe pump, connector tubing and a nozzle/nozzle holder combination. This can be seen in Fig. 1. Many of these components can be 3D printed and it is designed such that it can feasibly be attached to a wide range of desktop FDM 3D printers, as the only component in contact with the printer itself is the nozzle holder. We utilise the Creality Ender 3 as a low-cost and robust FDM platform, capable of running the material extruder.

**Figure 1:**
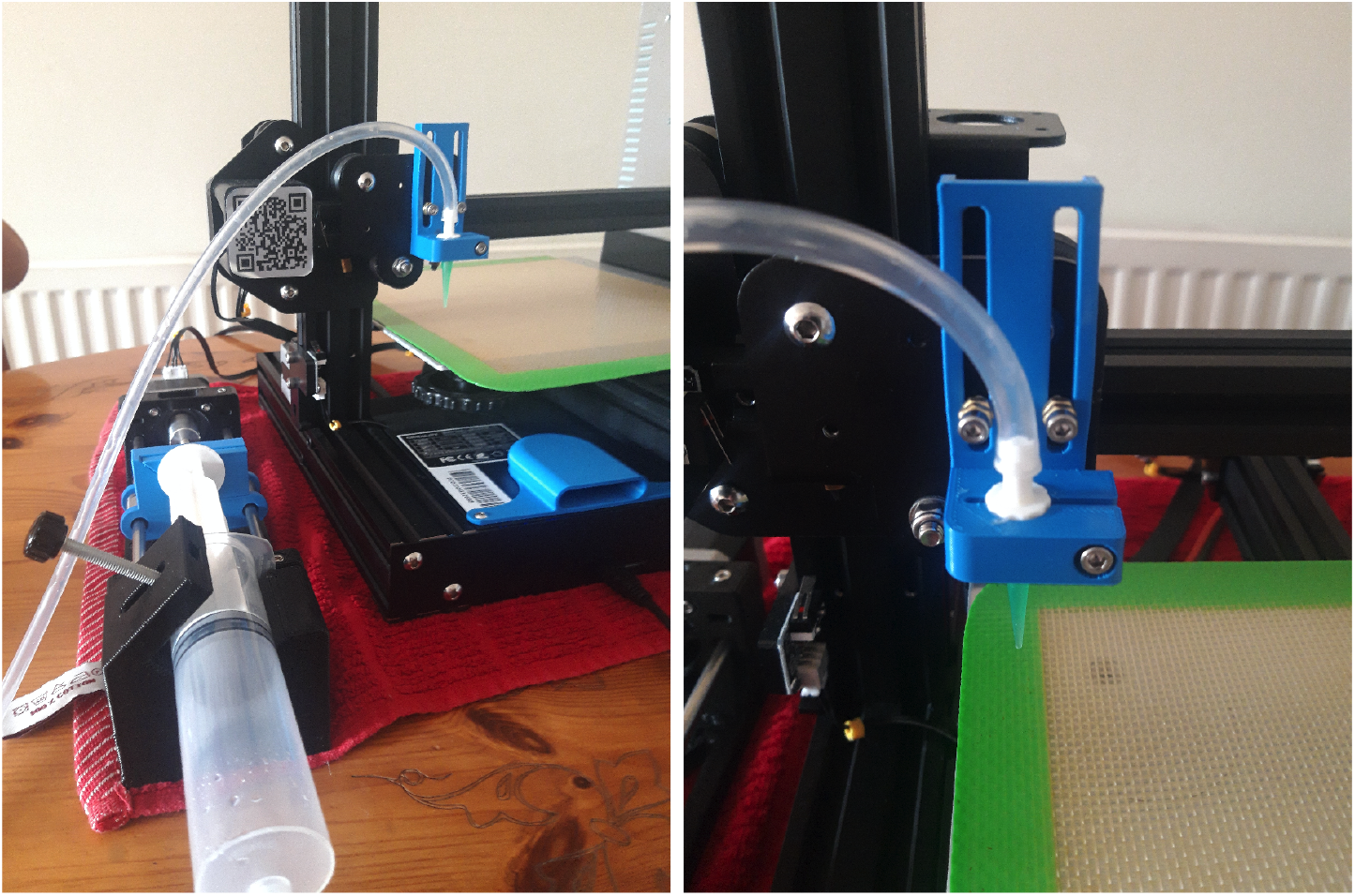
(a) Full paste extruder setup, including the syringe pump, connectors and nozzle, connected to Ender 3 3D printer. The plastic sheeting on the print plate is simply a non-stick baking sheet, for ease of removal of the print. (b) Nozzle holder design close up. Attaches directly the printing plate that holds printhead of the original 3D printer. The central holder design can be easily adjusted to accommodate a variety of connectors and tubing sizes.

**Figure 2:**
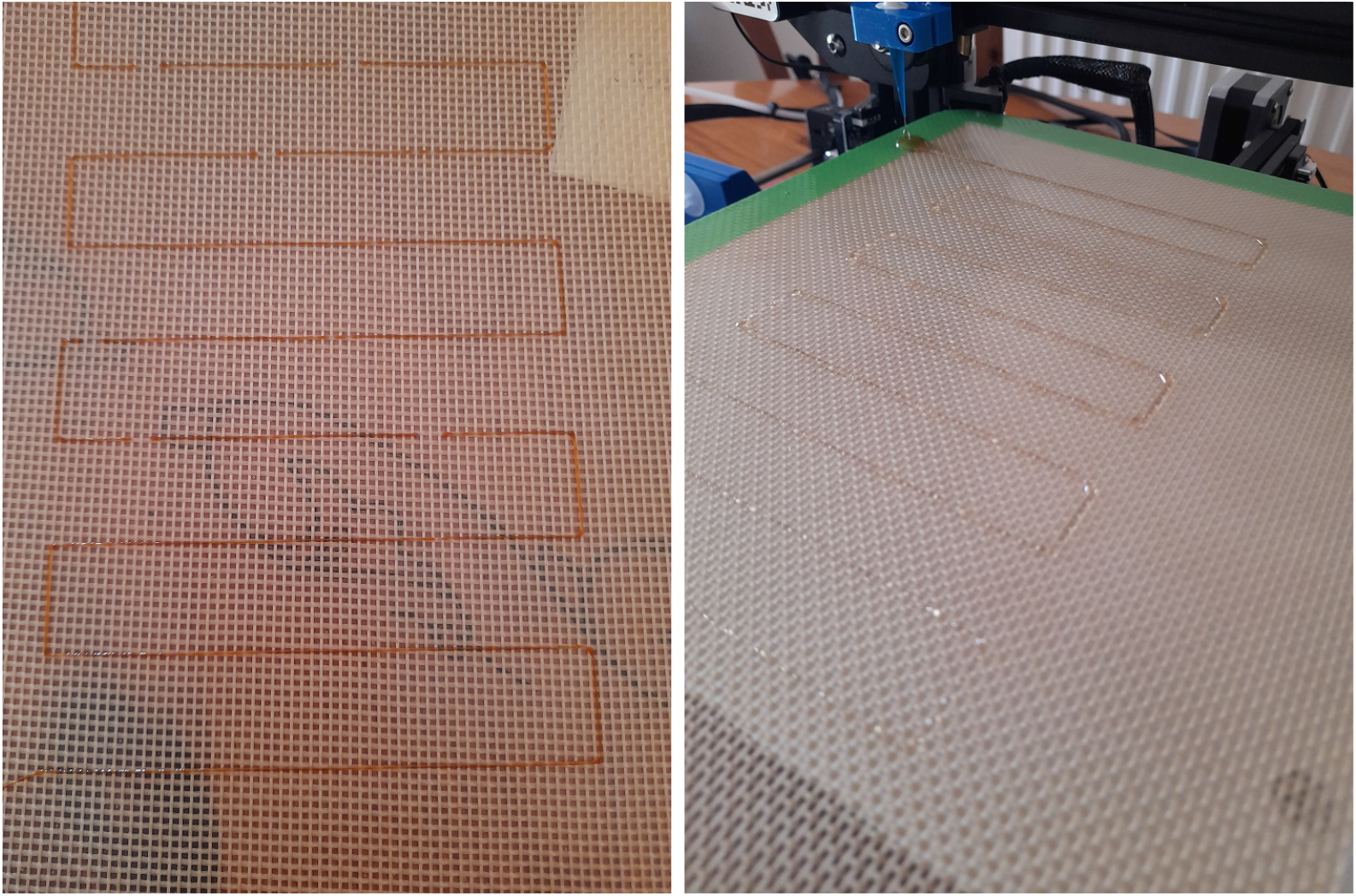
Examples of the low-lying scaffold print, varying the material (marmite and chitosan, respectively), the nozzle (0.5 and 0.75 mm, respectively) and the extrusion per mm but keeping the pattern the same.

Other printers may require modifications to the nozzle holder design or additional parts in order to work effectively. For instance, due to the choice of the Ender 3, we also recommend an additional 3D printed fan cover over the electronics housing to preclude the possibility of liquid inks leaving the print bed and entering the electronics housing. This may not be necessary on all printer types, however it is highly recommended that you search for any possible weakness points, prior to using the paste extruder.

Another useful, but non-essential, component to have is a sacrificial or non-stick surface on which to print. This makes removal of the print from the print bed far easier, reduces the amount of contact the original platform has with these paste inks and therefore increases the likelihood the printer could returned back to a traditional 3D printer state.

### 2.1 Syringe Pump

The ink reservoir and pumping system is a simple syringe pump, based on the impressive Poseidon Pump design [32]. This uses a stepper motor to drive a compressive wedge that acts on the syringe plunger. The body and driving wedge are both 3D printed, whilst the rest of the design is comprised of widespread components, found in both general hardware sellers and more specialist 3D printing storefronts. The wedge contains both either a flat surface or a small lip, upon which the syringe flange can be locked, in order to provide some amount of retraction to the syringe when driven in reverse. In addition, it provides an adjustable clasp so that a variety of syringe types and sizes can be used as the ink reservoir. In the original Poseidon design, the pump is driven by a Raspberry Pi/Arduino board, however here we instead power it via the desktop 3D printer using the extruder stepper motor JST connector, meaning any standard G-Code instructions can be interpreted by the paste extruder.

The pump is easily assembled, with both video and written instructions, and a low-cost bill of materials [32]. Alongside this, it is robust across multiple printing precision and layer heights, with only minimal finishing required to ensure all screw and bolt combinations fit. Finally, the ease of accessibility to the stepper motor means that it can easily be swapped out for other motors with additional torque or functionality. Unfortunately, the size of the original Poseidon Pump design is too large to be printed on the Creality Ender 3, however the design was adjusted by decreasing the height of the pump and therefore the separation between the stepper motor and the syringe holder and scaling the threaded and un-threaded rod required for assembly. This adjusted design is available on request from the corresponding author.

Other similarly produced syringe pump designs exist [33, 34, 35], alongside many commercial options, that could all easily be substituted in place of the Poseidon Pump.

### 2.2 Connector Tubing & Nozzle

The ink reservoir interfaces with the nozzle via male Luer lock connectors and plastic tubing. This locking system promotes even pressure during the extrusion process, but is not strictly necessary unless printing with at high pressures. A variety of sizing exists, however we find tubing of 5 mm in diameter has a simple interface with many common syringe sizes and types. Whilst a seemingly low impact component of the design, this can have a large effect on the printability of an ink and the pressure required for extrusion. Ideally, the distance between the reservoir and the nozzle should be minimised to minimise the pressure required for extrusion. However. to ensure that the nozzle can cover the full dimensions of the printer, a minimum length is required. The inner diameter however can be varied and can therefore be made to accommodate a variety of particulate sizes and viscosities -a seemingly unprintable mixture can be made more accessible by shortening the distance between reservoir and nozzle and increasing the inner diameter. The geometry of the extrusion apparatus has long been known to affect the pressure felt by the extruder and the material and can therefore be a route to both increasing printability and further affecting the material properties during extrusion [36, 37, 38].

The nozzle portion comprises again of a male Luer lock connector, joining the tubing and nozzle and held in place by a 3D printed holder, included in the Supplementary Information. The holder can be raised and lowered easily to ensure a variety of nozzles can be accommodated and provides a straight-forward route for adjusting the relative z-coordinate, necessary when printing directly onto a raised substrate. Due to the generic Luer lock connection, a variety of nozzles can be attached, with both plastic and metallic director components.

## 3 General Ink Formulation

As with the software component, the formulation component of the design can vary depending on the material used and the needs of the user. Generally, the ink can be thought of as a mixture of the desired components and any additives that ensure printability. Additives could be materials that affect the viscosity, the material strength or the surface tension -however care has to be taken that these improvements aren’t at the cost of other factors that impact the printability. The desired components consist of the core material to be printed and any other functional additives -for example pigments or other optically functional components. These components can also be processed to ensure more practical printability, including reducing particle sizes or chemically modifying the material. More often, care has to be taken in modifying these core materials to ensure that the final printed product retains the desired properties.

**Table 1:** Some examples of printable ink formulations and the parameters varied in order to enable their printing. Note that all of these examples use differing printing equipment and may be over-simplifications of the full process this list is intended to highlight explicitly mentioned commonalities in parameters that increase printability and the types of materials that can be printed.

| Ref. | Core Material | Additives | Additional Parameters |
| --- | --- | --- | --- |
| [39] | Chocolate | None | Temperature control<br>(heating in reservoir,<br>cooling on bed) |
| [40] | Marmite/<br>Vegemite | None | Temperature control<br>(heating in reservoir) |
| [41] | Mashed<br>Potato | Potato Starch | None |
| [42] | Chitosan | Acetic Acid<br>& Raffinose | Temperature control<br>(cooling on bed) |
| [43] | Chitosan | Hydroxyapatite,<br>Acetic Acid,<br>Glycerol Phosphate<br>Disodium Salt<br>& NaOH | None |
| [44] | Alginate | Sodium<br>Periodate | Temperature control<br>(cooling on bed) |
| [45] | Kaolinite | Water | None |
| [46] | Poly-<br>(ε-capro-<br>lactone) | None | Temperature control<br>(heating at nozzle)<br>& slow print speed |
| [47] | Concrete | Super-<br>plasticizer,<br>retarder &<br>accelerator | Particle size chosen to<br>match nozzle size |

External parameters can have a direct impact on the printability as well, including ambient airflow, pressure applied and the temperature of the ink, surroundings and printbed (the interplay between these three can be quite complex for certain materials).

## 4 Print Tests

Traditional tests of 3D printing platforms typically involve a complex design such as Benchy [23], in order to showcase the level of detail and complexity possible. For paste extruder designs and viscous inks, a differing judgement of the overall quality must be made, that are dependent on criteria defined more by the desired output. However, we can make generic statements about the properties of the print using these tests that can then be used to further fine tune the printing parameters and improve the overall resolution, repeatability and printability.

### 4.1 Low-Lying Scaffold Design

The first test involves printing a simple line design that incorporates multiple 90° turns and a handful of layers, similar to the scaffold designs that are widely used in bioprinting [24, 26, 28]. This is designed to probe three main properties of the print:

- The general thickness of a printed single line, with respect to the nozzle diameter.
- Broadening of a 90° bend.
- Possibility and common issues of layering.

An example of G-Code used to produce a design of this type has form

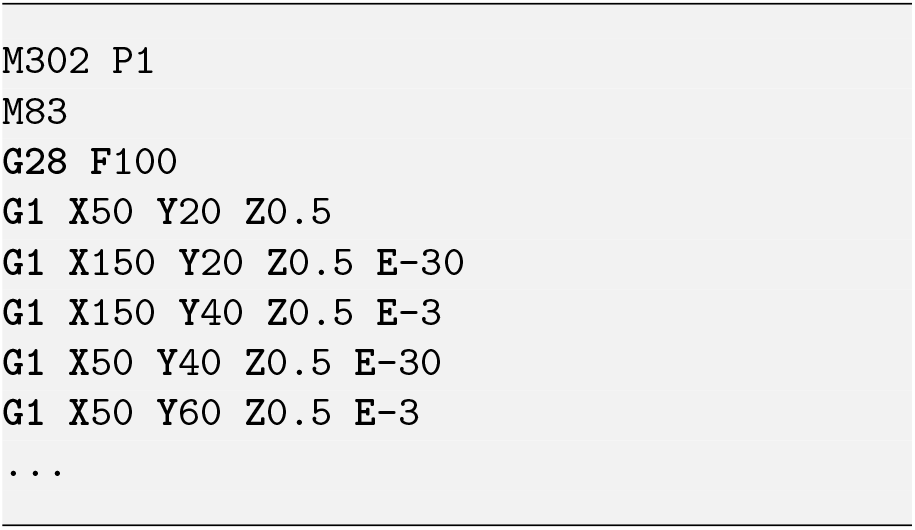

This can repeat giving multiple horizontal sections to the design and there-for a long observed print time and an increased chance of successful print. The opening *M* statements allow for cold extrusion and sets the extrusion coordinates to be relative, as opposed to absolute. This is particularly important for the syringe extruder where there is no longer a definitive zero, as this depends on the filling of the syringe reservoir and the amount of material available. The *G*1 commands are direct movements with (*X, Y, Z*) coordinates and an extrusion command *E* - the value following this command can initially be modified by hand so that it that it better aligns with the material response of the ink under pressure, the stepper motor in use and the geometry of the ink reservoir and connectors. This allows for a rough estimations of the required extrusion per mm steps.

Sample variations of this code are included in the Supplementary Information, for a variety of layer numbers. An additional Python script is included that takes in a coordinate set defining the maximal XY values of the scaffold, a extrusion multiplier and the number of 180 degree turns to incorporate per layer. In this setup, each subsequent layer is rotated 90 degrees to the layer beneath.

The thickness of a single printed line, typically the horizontal sections, can be quantified as a function of the nozzle diameter as

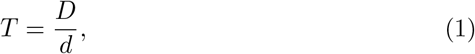

where *D* is the thickness of the printed line and *d* is the nozzle diameter. It is best to obtain multiple measurements of *D* and average across them. Additionally, this value may change (sometimes dramatically) with increasing number of vertical layers, due to the flow and adhesion properties of the ink. As such, we define a complimentary value *H*, linked to the expected and measured overall height

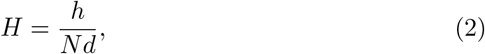

where *N* is simply the number of layers attempted. A combination of *T* and *H* values can be used to determine the flow of the ink post-deposition and therefore the feasibility of verticality for this particular formulation. Note however that even in standard 3D printing systems and materials such as PLA, this value does not tend towards one. Standard values for PLA are in the region of *H* = 0.8.

Building from this, the broadening at the corners can be defined as

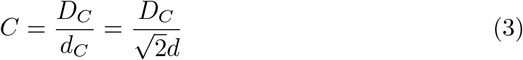

where *d*_*C*_ is the expected corner length, based on the nozzle diameter, and *D*_*C*_ is the measured corner length.

Note that for the same G-code these values will vary with the size of the ink reservoir, the ink formulation and, to a lesser extent, the connection between the syringe pump and nozzle -therefore it is important to keep these parameters constant during data collection.

**Table 2:** Scaffold design parameters and routes for tuning the print properties from these tests.

| Print Property | Problem | Possible Cause | Possible Solution |
| --- | --- | --- | --- |
| T | $T \gg 1$ | Overextrusion OR Die Swelling | Increase nozzle diameter (may be limiting factor to print resolution) OR decrease extrusion per mm |
| T | $T \ll 1$ | Underextrusion | Increase extrusion per mm |
| T | T varies noticeably across print | Uneven extrusion | Increase nozzle diameter OR decrease particulate size |
| T | $T \approx 0$ | Clogging | Increase nozzle diameter OR decrease particulate size OR degas ink |
| T, H | $H \ll 1$<br>$T \gg 1$ | Flow after extrusion | Increase viscosity of ink OR decrease temperature |
| C | $C \gg 1$ | Pooling and Overextrusion | Retraction & Coasting |
| C | $C \ll 1$ | Underextrusion | Too much retraction |

### 4.2 Slump Test

Similar to the concrete slump test [48], we attempt to print a conical 3D object in order to further characterise the readiness and issues with a particular ink formulation in vertical layering. Traditionally, this test would involve a concrete mixture that would be gathered in a conical mold, with the mold removed carefully, to examine the loss of verticality or the ‘slump’, due to the competition between gravity and the setting of the mixture.

Here, we can build a series of conical shapes of differing heights and widths to examine both the slump and the frequency and types of errors in the stacking. These are most easily built in TinkerCAD, using the in-built cone feature, and can quickly produce a variety of designs through varying the height and

We define both the ratio between the diameter of the cone at the base and top and their expected values,

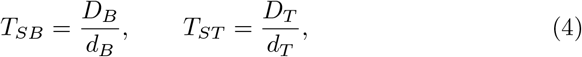

where a subscript *B* represents a measurement at the base, a subscript *T* represents a measurement at the top of the cone and *d* represents the expected diameter at those points. Depending on the amount of the frequency of printing faults occurring, these values may not be rotationally symmetric and therefore an average across multiple angles may provide the most accurate representation. Additionally, depending on the material used, an accurate measurement of the top width may be impossible due to varied levels of slump around the print, as such we use the uppermost height of the cone as a third comparative term

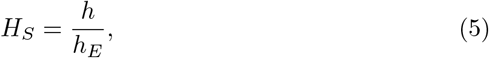

where *h*_*E*_ is the expected value of the cone height. However, as with the scaffold design, the value of this ratio tends toward 0.8 for a traditional 3D printing materials such as PLA.

Variations in the height value are also useful for tracking any time-dependent properties of the printing ink, such as setting times and the separation of emulsions. Materials under a prolonged duration of pressure, such as those in the reservoir, can respond differently, particularly in terms of mixing and setting [49, 50], and produce a differing response in terms of the printed line width and height.

### 4.3 Step-Wedge Print

Other print features that are present and provide useful information about the print properties are more difficult to quantify. As such, a generic three-dimensional printed structure is useful for determining the suitability of an ink for more vertical prints, the interactions between differing types of print feature (walls/perimeters, infill, surface texture etc.) and other properties of the printer and slicer combination, like the print speed and rate of extrusion.

For this, we use a simple step-wedge design consisting of 10 adjacent steps, ranging from 1 mm to 10 mm in intervals of 1 mm. Each step is 1 cm x 1 cm in the XY dimension and provides a chance for printing both walls and infill at each layer and probing interaction formed by both vertically and horizontally neighbouring them. Note that the infill is always chosen to be 100%, simply because the vast majority of materials are unable to self-support themselves without the presence of material in the layer beneath.

These issues can be hard to troubleshoot, however generic print defects such as inconsistent wall thicknesses and overhangs can be reduced most commonly by reducing the printing speed and increasing the number of perimeters printed. Stringing is other common problem that can instead be mitigated by reducing the ‘travel speed’, the speed at which the nozzle moves when not printing. These problems are complex combinations of many parameters and may therefore not be possible to eliminate entirely.

Similar height and width measurements can be used for this print in order to determine the overall effects of setting and flow after deposition, however the previously described tests are more efficient in terms of time and material usage.

## 5 Case Studies

### 5.1 Gelatine

Gelatine represents an interesting test case, due to the prevalence of the material within bioprinting, most prominently as gelatine methacrylate (GelMA) [5, 51] or a gelatine/alginate composite [5, 52], as opposed to pure gelatine. These materials are favoured due to their enhanced mechanical properties via induced crosslinking through, for instance, the addition of UV irradiation and calcium chloride respectively [5]. Gelatine alone also physically crosslinks or sets upon cooling to room temperature [53] and the rate of setting is directly linked to the concentration and the Bloom of the sample (a measure of the setting strength).

For a purely gelatine and water ink, 10-20wt% mixes of 250 Bloom gelatine heated to 40 -50°c give a window of feasibility small enough to set once deposited but large enough to retain flow during the majority of printing. This attempts to balance the structure provided by the fast setting rate and the rate of extrusion, so that the material sets on the build-plate rather than in the reservoir. Failure of this system usually occurs due a mismatch of these competing effects leading to setting of the ink within the nozzle and reservoir, however we find the above parameters allow for volumes in the region of 10 -20 ml of ink (or 5 -10 print layers, depending on the design) to be extruded. Examples of gelatine prints can be seen in Fig. 3, including high-resolution and layered lines, filled two-dimensional shapes, a layered slump test and a smoothly increasing stepped system.

**Figure 3:**
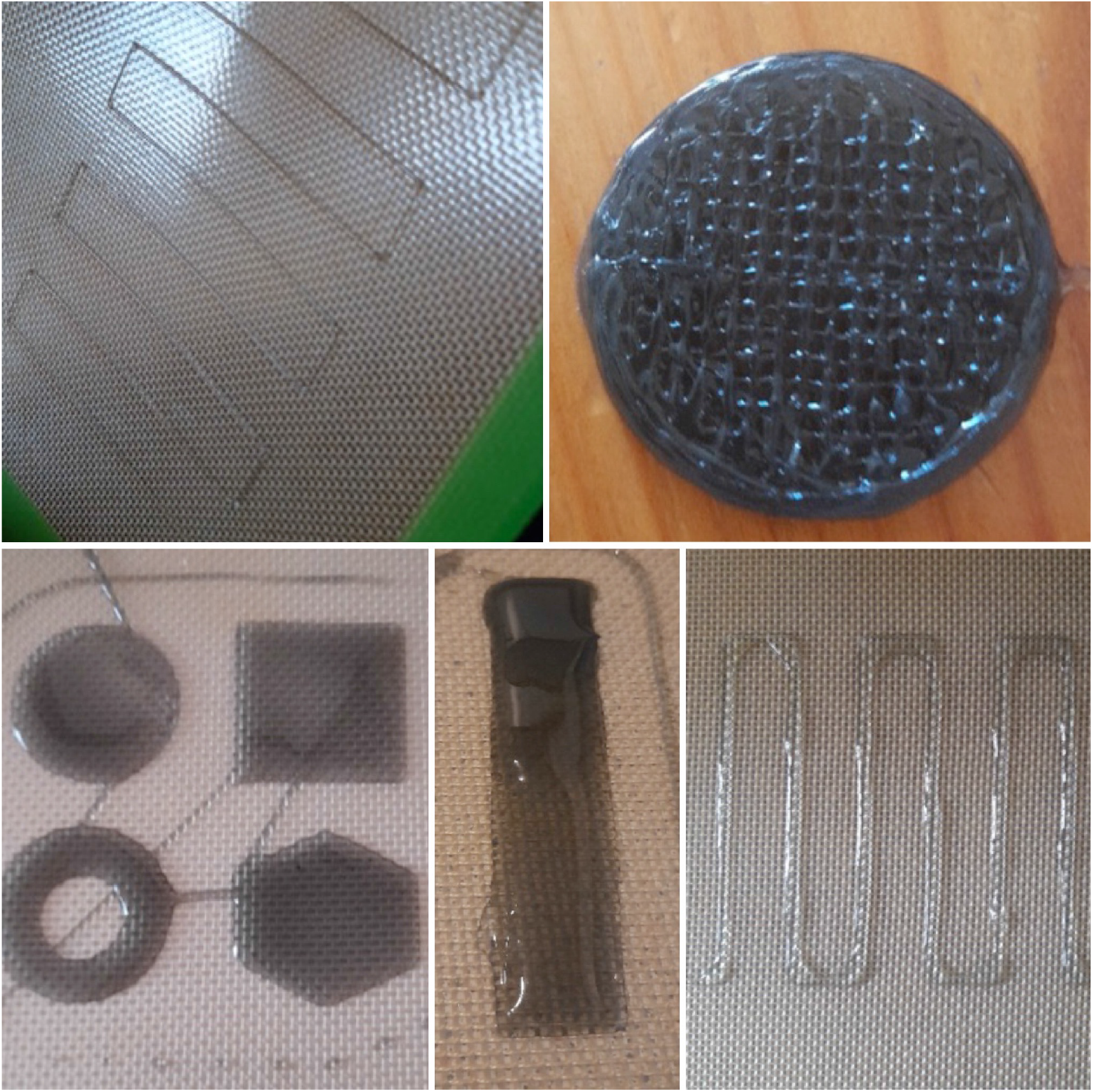
Collection of structures printed with gelatine, including thick (3 mm) and thin (1 mm) single lines, slump tests and low-lying vertical structures like the step-wedge. In each case, the ink consists of a 10-20 wt% mix of gelatine, water and a small amount of carbon black pigment, for visual clarity. This highlights the common drawbacks of printing the material, including loss of detail due to flow, retraction issues and setting during the printing process.

Due to the effectively liquid state in which the ink leaves the nozzle, the resolution of a single line is limited by a combination of liquid flow on the printbed and die swelling effects. Measured line widths are regularly in the region of 1 -2 mm, with a nozzle width range of 0.4 -1.7 mm, and right-angled corners display a similar and more pronounced broadening effect. The data of which can be found in Fig. 4. Additionally, due to the slowly increasing viscosity provided by the lack of temperature control, this line width appears to decrease over time, ensuring the slump test succeeds up to as many as ten layers (circular print in Fig. 4) and overhanging layers are possible for small overhangs (rectangular ramp increasing in height and overhang).

**Figure 4:**
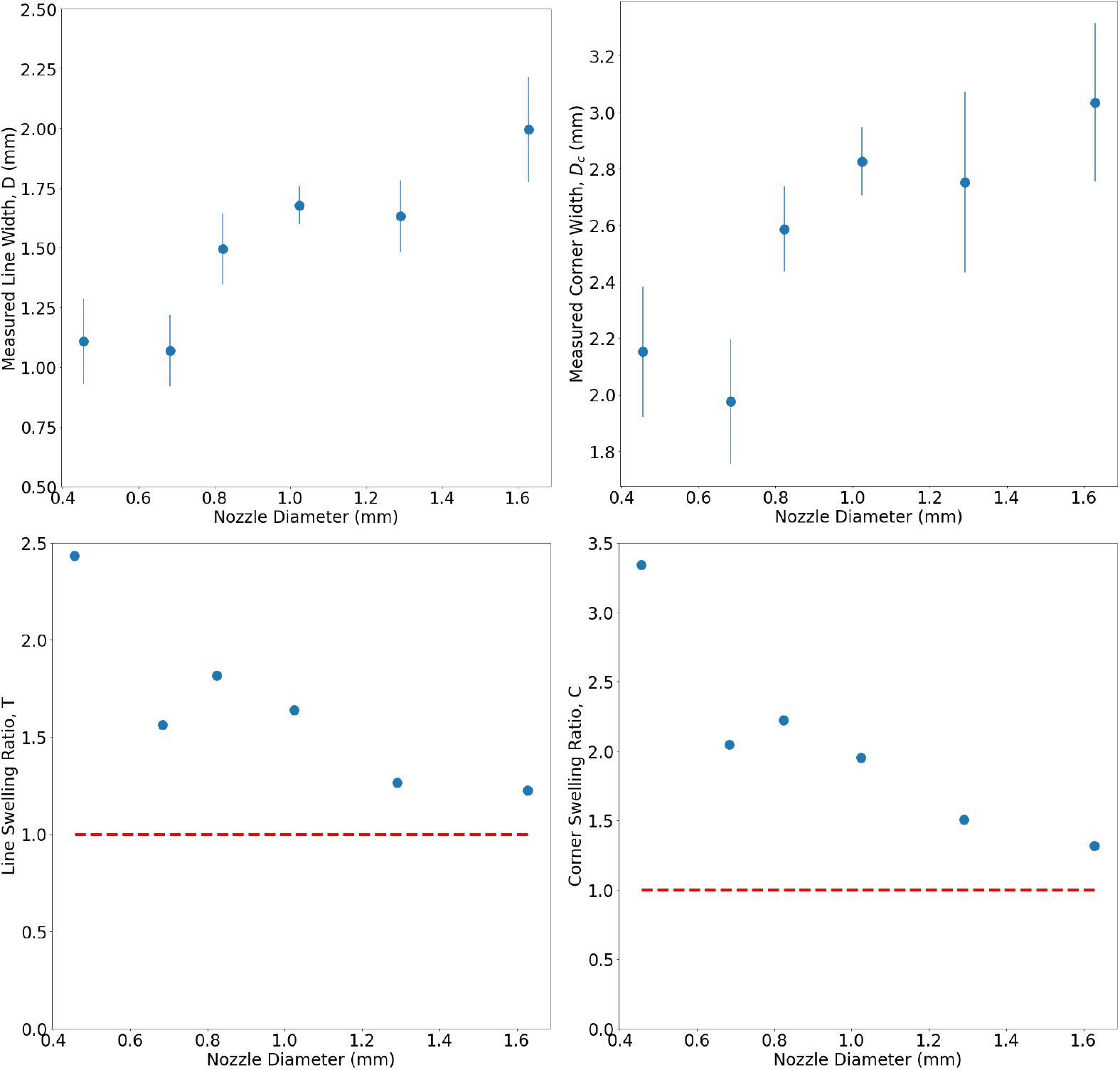
(Above) Data for the measured line and corner width of a 17.5wt% aqueous gelatine mix held at 40°c and averaged over three measurements. (Below) Data for the swelling of the individual line and 90° for printed gelatine at a variety of nozzle widths. As expected, the swelling ratio decreases as the nozzle width increases.

This particular ink and hardware combination could be improved by focusing on the temperature effects across the extrusion. Setting within the reservoir and nozzle could be avoided by heating within these sections to reasonable temperatures of 40 -50°. This could be combined with a cooling of the printbed in order to discourage flow of the ink upon leaving the nozzle. Alternatively, an ink formulation driven approach could seek to use a higher concentration of gelatin or a gelatin component with a higher Bloom, in order to encourage quicker setting on the buildplate.

The slump test for gelatine, seen in Fig. 3, produced useful data beyond the simple dimensional data. It was found that as the print was undertaken, the printed line width decreased until the ink mixture set in the nozzle and the print was no longer achievable. For a mixture heated to 40°c, measurements of the line width were taken from a continuous line print, noting the time from initial extrusion. The continuous printing, with no periods of retraction, encourages a longer suitable print window before clogging. This is shown in Fig. 5, displaying a near halving of the print width over the course of a 10 minute print.

**Figure 5:**
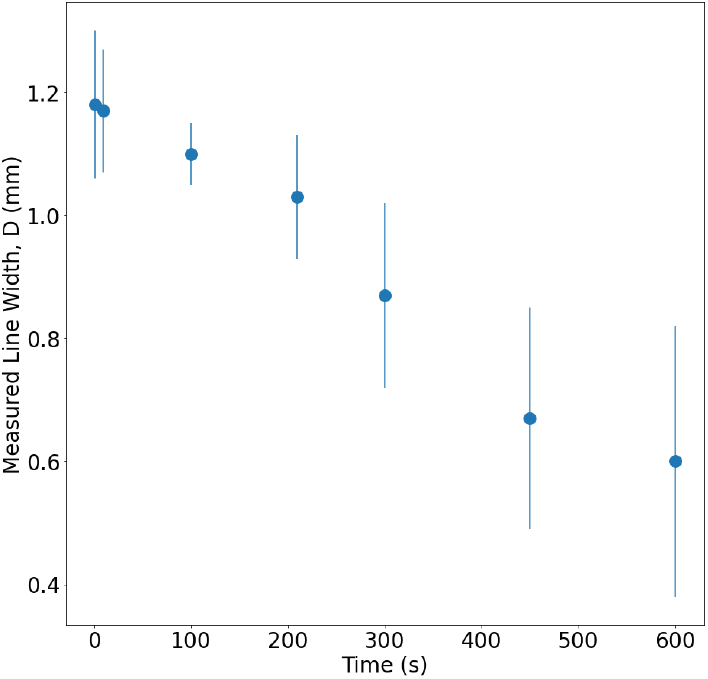
Data showing the decrease in linewidth with time, due to the setting of the gelatine ink within the reservoir and nozzle. Each data point is averaged over three attempted prints. Beyond 600 seconds, the ink had set too much to be extruded.

Conical sections up to 1.5 mm were produced before the setting stalled the print, giving a height shrinkage value of *H*_*S*_ = 0.65 and ratios of the bottom and top diameters as *T*_*SB*_ = 1.24 and *T*_*ST*_ = 0.92 respectively. This suggests a flow after deposition on the build plate, that expands the lower diameter and decreases the upper diameter. It should be noted, however, that the upper diameter (that should be well defined due to the shape of the input 3D model) was instead hard to identify in certain places in the print due to the ink flow and so measurements were hard to take.

Finally, the step-wedge print test, also seen in Fig. 3, was produced. This is useful for highlighting the generic issues with the printing material, including the loss of detail due to material flow, retraction values being hard to control and decrease in the printed line width with time due to setting of the ink.

### 5.2 Chitosan

Chitosan is another similar material to gelatine, providing high biocompatibility with a non-toxic and inexpensive material [42]. Unlike gelatin however, it has a long gelation time and therefore is immediately a less suitable material for printing. Despite this, it provides a useful vector for probing how the movement of the ink on the printing plate after deposition.

A mixture of 1:9 of chitosan powder and household vinegar (in place of acetic acid) was produced by mixing by hand and leaving for 24 hours in a cooled enviroment. This allowed for the complete dissolution of the chitosan powder and an extrudable but viscous solution. A simple scaffold design was printed, one layer high, and the line width measurements were now monitored over time -the results shown in Fig. 6. As expected, the slower gelation times resulted in a slow dilution of the print line, increasing the measured width, and explaining the common need to alter chitosan-based inks with other additives to make them more suitable for printing [42, 43].

**Figure 6:**
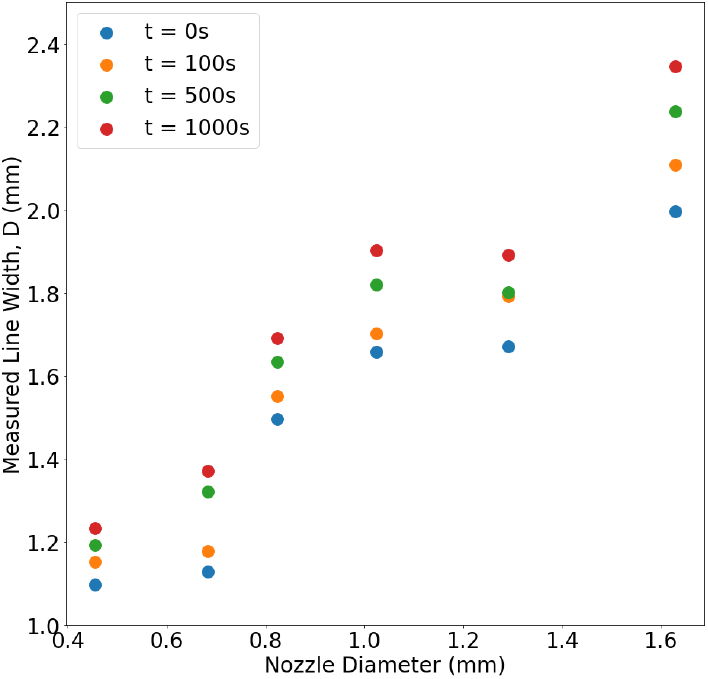
Data showing the increase in linewidth with time, due to the slow gelation time of chitosan. Each data point is average over three attempts. Beyond 1000 seconds, marginal differences were seen in the line width measurements, however the ink had only partially set.

### 5.3 Marmite

Marmite presents a simple case for a conductive paste, in order to produce custom (and edible to some) circuitry [40]. It is a highly viscous material that exhibits shear-thinning behaviour, which increases with increasing temperature. Despite this, it has been shown to be readily printable at room temperature using a BioBot 1 extrusion printer [40]. We use this test case to explore how a functional viscous material may be extruded, without the strict temperature dependence and setting behaviour exhibited by a material such as gelatine.

Initial extrusion tests show that the highly viscous behaviour of the material means that in order for consistent extrusion to take place, the feedrate must be slowed and the delay between the stepper motor instructions and deposition from the nozzle mean that retraction is an inherent issue. We find however that once the ink is moving under inertia, the printed single lines are of a consistently high resolution, hindered only by the presence of air introduced by placing the highly viscous material in the syringe.

As there is no setting component to the ink, we can assume that anything we print will slowly flow, regardless of structure, and so aim only to explore the range of designs that are relatively fast, on the scale of minutes. A key component of the work in Ref. [40, 54] is the production of electrically conductive channels -slower printing with larger nozzle diameters can produce thick conductive routes on a variety of substrates, as seen in Fig. 7.

**Figure 7:**
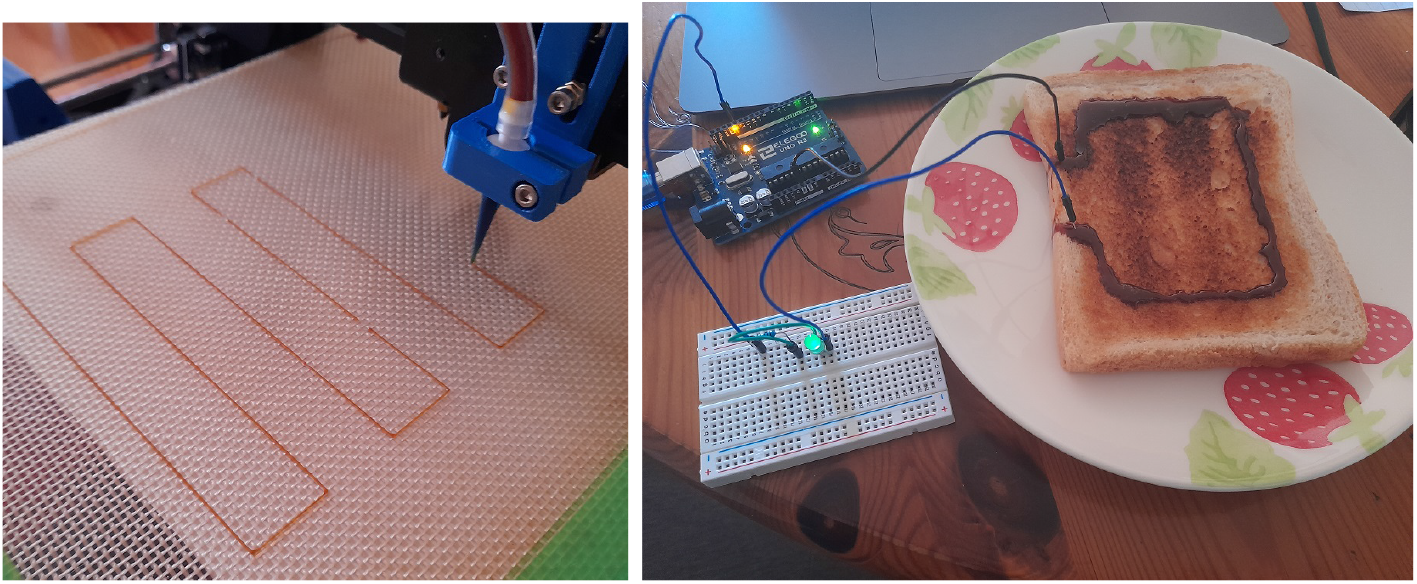
Using Marmite as an ink within the paste extruder system. No changes are made to the material and it is printed at room temperature, showing thin printed lines that are affected by the presence of air bubbles in the system (Left). When printed at slower speeds with larger nozzles sizes, these dislocations become less important and thick conductive lines can be printed to make similar circuitry to that in Ref. [40]

**Figure 8:**
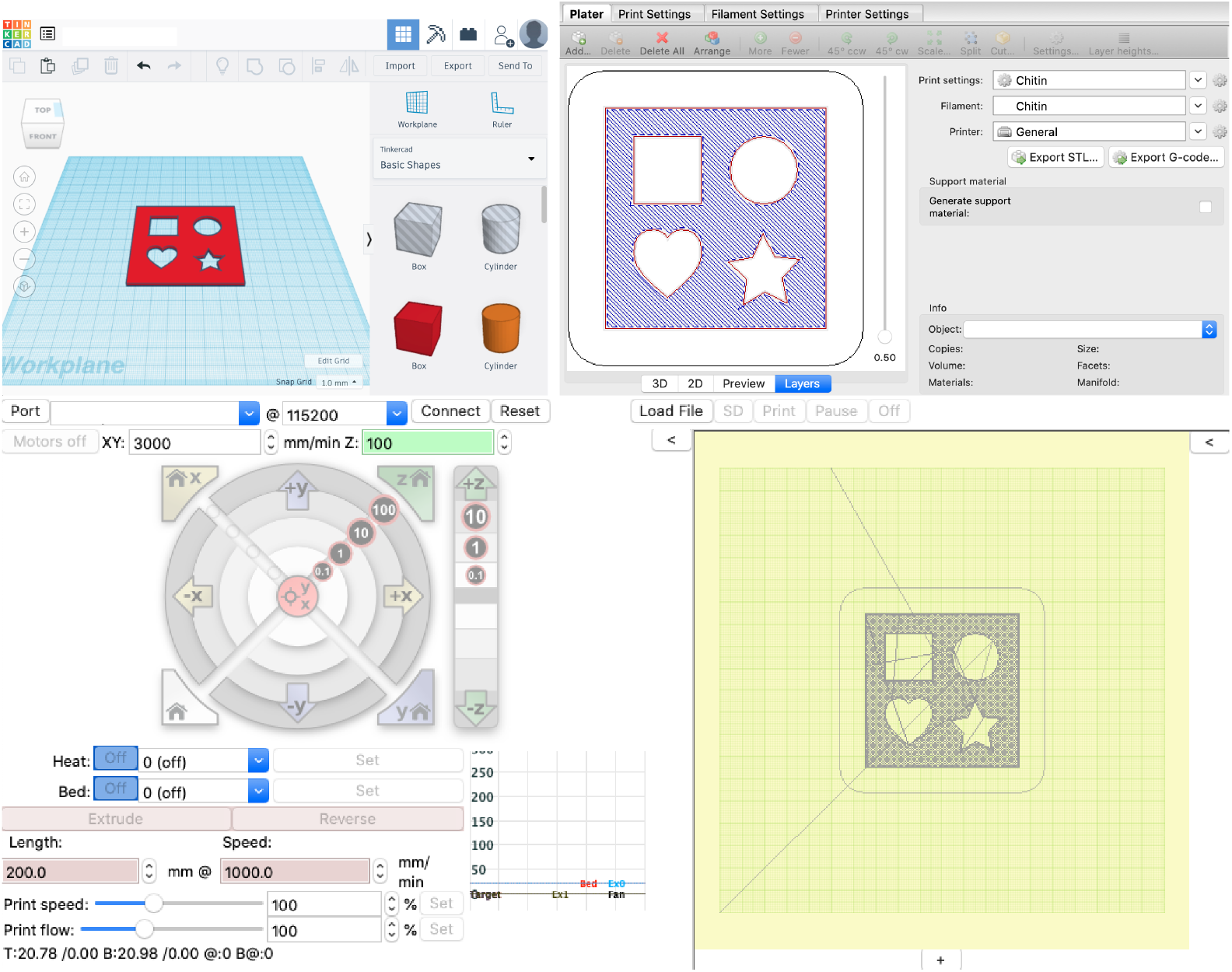
Examples of the software component of the paste extrusion system. (Upper left) A CAD programme, for design of 2D and 3D objects for printing. Here we highlight TinkerCAD, a free browser based CAD programme, useful for beginners. (Upper right) Slicing software, in this case Slic3r, for producing custom ink-specific instructions per object. (Lower) Printer interface for monitoring and sending the instructions produced by the slicing software. All together this shows one possible workflow for the software component of the paste extruder setup.

## 6 Conclusion

We present a low-cost and easy-to-assemble platform for the extrusion and selective deposition of gels and pastes, when combined with a generic desktop 3D printer. This is designed to be easy to use and manipulate, in order to make it compatible with a wide range of printers and printing material. As such, we utilise a handful of extremely simple printing tests that can be used to produce material profiles, to help move toward repeatable and clean prints. Using these, we explore a handful of material case studies, differing in viscosity and setting properties, and highlight how early testing data could be used to produce specific material/printer profiles.

## Appendix A Software

The software component of the system comprises of three distinct responsibilities; a Computer Aided Design (CAD) component, a slicer and an interface with the printer. Certain aspects of these may be integrated into one software package, but it can be useful for the purposes of the workflow to have them separated. The CAD component is present to provide a work space for the design of 2D/3D objects for the printer. There are a huge range of options when choosing CAD software, each with varying focuses and degrees of complexity, but with many overlapping premises and approaches, so much of it comes down to personal preference. The slicer component is used to produce profiles that determine exactly how a desired three dimensional object will be broken into two-dimensional slices and then translated into the G-Code instructions for the print. These profiles can be tuned to be material specific, based on terms including, but not limited to:

- Printed line width
- Layer height
- Extrusion per mm
- Print speed
- Infill type and properties
- Perimeter settings of external walls
- Brim loops

For this, we recommend Slic3r [55] and provide a collection of material profiles that have been tested on the paste extruder setup within the Supplementary Information. These can be used as starting points for similar materials, however they may require further tuning for each particular ink formulation and printer combination.

For the interface, we use either Pronterface [56] or Octoprint [57]. Both allow for line-by-line live entry of G-Code and therefore can highlight the particular instructions and movements that may be causing failure within the print. More generally, any interface that allows for monitoring of the print, live pausing and resuming and the sending of instructions directly should be suitable.

## Appendix B Standard Procedure

A general workflow for the extrusion of a generic ink mixture may look something like:

- **Digital Design** -Production of desired STL in CAD design software (Fusion 360, Rhino, Tinkercad etc.) and translation to G-Code via slicing software.
- **Ink Formulation** -Identification of desired printing material and the routes of processing and modification required to produce a sufficient ink.
- **Slicing Parameters** -Variation of the slicing parameters in slicing software to ensure that the interpretation of the digital model matches the limitations of the ink. This stage is the most obvious point of variation for iteratively finding a combination of software and material properties that ensure a printable ink.
- **Reservoir Filling** -Syringe must be filled with the chosen ink recipe. For particularly viscous inks, this can cause issues with trapping air pockets that can be overcome by vacuum degassing or simply allowing time for the air bubbles to rise to the surface. Air bubbles will cause intermittent extrusion issues and variation in the pressure applied to the ink by the syringe pump.
- **Priming Syringe Pump** -Due to the relative nature of the extruder stepper motor coordinates, it often defaults to a zero position upon a reset of the printer, no matter the actual position of the compressive wedge. It is theoretically possible to change these settings within the printer, however this limits the rate at which the machine can be returned to a simple thermoplastic extruder and so we choose to work around it. It is difficult to determine a set of G-Code instructions that will ‘home’ the extruder motor and we instead drive the stepper motor manually to the desired starting position, using linear move (G1) commands. For example: 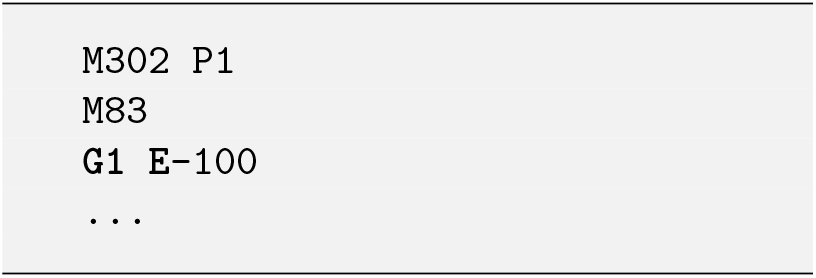
- **Loading Syringe Pump** -Once the syringe pump and filled syringe are aligned, the syringe is loaded into the pump and secured with the cradle screw. The nozzle and the printbed are aligned by hand, typically with the same tolerance you would aim for in a standard 3D print levelling process. Extrusion commands are sent manually to the machine until the connector tubing and nozzle and filled with ink.
- **Build Area Test** -Interaction between the syringe pump and printer dimensions are tested, including ensuring the connector tubing does not interfere with the movement of the print. Below is a series of generic instructions that can be used to probe this movement, the magnitudes may vary between printer types. 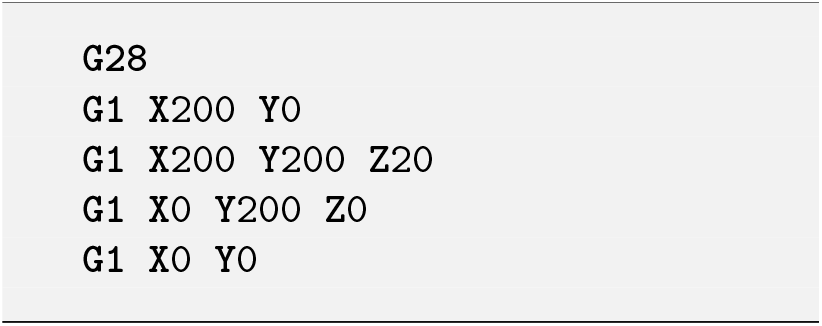
- **Extrusion Test** -To ensure there are no regions of pressure build-up and potential failure, a short extrusion test should be undertaken prior to printing. This is another chance to ensure the nozzle tip and printbed are sufficiently spaced. The code below moves to a uncommonly used space, initiating a small extrude move, and then tests the extrusion at the plate and above the plate, before retracting a little to ensure any induced flow slows down. 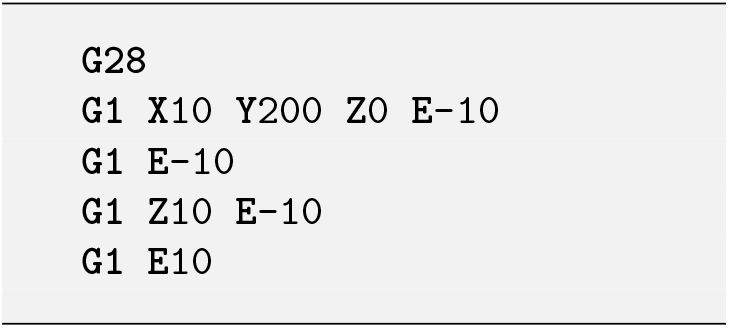
- **Running Print** -Finally, the print instructions can be sent to the printer. The printer should be monitored during this phase to ensure there is no clogging or stepper motor skipping that could damage the print setup.

## Appendix C Inherent Advantages & Limitations

This extruder design has some inherent advantages over commercial paste extruders and bioprinters, such as:

- A low price. Components are either 3D printed or comprised of commonly found and cheap items. The majority of the cost arises from the original 3D printer.
- Compatibility with a variety of desktop FDM 3D printers. This is due to the majority of the design being external to the printer, with only the nozzle portion being in contact the 3D printer.
- The design is modular, allowing for individual components to be swapped depending on the users needs. Common variables include the syringe, nozzle and tubing dimensions and the choice of stepper motor. Additionally, this design could also incorporate a range of further external modifiers, including temperature control and optical interactions.
- Can easily be converted back into a plastic 3D printer.
- Wide range of material choice. Modification to both hardware and software is always an option to increase printability.
- Requires little specialist knowledge. The ease of assembly and intentionally straight-forward nature of the design make this accessible to a wide variety of academic backgrounds, as well as being a useful and visually impressive tool for education and outreach.

It also has some inherent limitations, the understanding of which can help with material choice and ink formulation. These include:

- Uneven extrusion due to plastic warping, present in both the body of the syringe pump and the syringe itself. This could be partially averted in the latter case through the use of a more structurally stable glass or metallic syringe.
- Retraction is limited, particularly in comparison to air pressure methods. This can induce overextrusion and stringing and make it difficult to fabricate objects where the print cannot be completed in a single continuous line.
- The reservoir size is large but still limited. Whilst this is suitable for low-lying prints of tens of layers, it cannot extend to the furthest reaches of the possible print dimensions. This is due to the design of the syringe pump and so could be side-stepped through modification of this stage.
- Connector tubing increases the chance of stepper motor skipping with increasing length. This can be avoided by minimising the length of the connector tubing and allow ample time for the system to settle between extrusion commands when initially filling the tubing.
- There is no failsafe implemented. As such, if an ink formulation is less reliable (or even unprintable), the design causes stress on the body of the syringe pump and the stepper motor by skipping steps. This is particularly true during larger prints, in which the printer orchestrates many instructions in succession and can cause a pressure build up faster than the material can be effectively extruded. In these case, a reduced feedrate can result in more even extrusion and less chance of stepper motor skipping.

